# Hippocampal subfields and their neocortical interactions during autobiographical memory

**DOI:** 10.1101/2022.05.03.490407

**Authors:** Pitshaporn Leelaarporn, Marshall A. Dalton, Rüdiger Stirnberg, Tony Stöcker, Annika Spottke, Anja Schneider, Cornelia McCormick

**Author notes:** Corresponding author: Cornelia McCormick, Pitshaporn Leelaarporn. Address: University Medical School, Department of Neurodegenerative Diseases and Geriatric Psychiatry, Venusberg-Campus 1 / 53127 Bonn, Germany.

## Abstract

Advances in ultra-high field 7 Tesla functional magnetic resonance imaging (7T fMRI) have provided unprecedented opportunities to gain insights into the neural underpinnings supporting human memory. The hippocampus, a heterogeneous brain structure comprising several subfields plays a central role during vivid re-experiencing of autobiographical memories (AM). However, due to technical limitations, how hippocampal subfields differentially support AM, whether this contribution is specific to one portion along the hippocampal long-axis, and how subfields are functionally connected with other brain regions typically associated with AM retrieval remains elusive. Here, we leveraged technical advances of parallel imaging and employed a submillimeter Echo Planar Imaging sequence over the whole brain while participants re-experienced vivid, detail-rich AM. We found that all hippocampal subfields along the long-axis were engaged during AM retrieval. Nonetheless, only the pre/parasubiculum within the anterior body of the hippocampus, contributed over and above to AM retrieval. Moreover, whole-brain functional connectivity analyses of the same data revealed that this part of the hippocampus was the only one that was strongly connected to other brain regions typically associated with AM, such as the ventromedial prefrontal cortex (vmPFC) and medial/lateral parietal regions. In the context of the broader literature, our results support recent proposals that the anterior body of the pre/parasubiculum may play an essential role in scene-based cognition, such as the re-experience of personal past events.

**Highlights:**

- All hippocampal subfields differentiate AM retrieval from mental arithmetic problem solving
- The anterior body of the pre/parasubiculum engages in AM more than other subfields
- The anterior body of the pre/parasubiculum is strongly connected to the AM network
- The pre/parasubiculum may be preferentially involved in scene-based cognition

## Introduction

Neuropsychological and functional magnetic resonance imaging (fMRI) studies have firmly established the hippocampus as a central structure underpinning vivid autobiographical memory (AM, i.e., memories of personal past events). The hippocampus is a heterogeneous brain structure comprising several subfields including the dentate gyrus (DG), cornu ammonis (CA) 1-4, subiculum, presubiculum, and parasubiculum (hereafter referred to collectively as the pre/parasubiculum). The hippocampus interacts with a broader set of brain areas that together comprise the AM network. This network includes areas in the ventromedial prefrontal cortex (vmPFC) and medial/lateral parietal cortices (Addis *et al*., 2007; McCormick *et al*., 2015, 2020; Moscovitch *et al*., 2005; Rosenbaum *et al*., 2008; Scoville and Milner, 1957). While we have a broad understanding that the hippocampus works within this network to support AM retrieval, we lack a detailed understanding of how hippocampal subfields interact with cortical areas of the AM network during AM retrieval. To address this gap, we leveraged recent advances in ultra-high field 7-Tesla functional magnetic resonance imaging (fMRI), to; (i) examine the contributions of hippocampal subfields to AM; (ii) assess how this differs along the anterior-posterior axis of hippocampal subfields and; (iii) characterize their associated functional connectivity with the neocortex.

AM retrieval is a complex cognitive process supported by a dynamic interplay between brain areas within the AM network (Conway and Pleydell-Pearce, 2000; Conway, 2009, McCormick et al. 2015). It is widely acknowledged that the hippocampus plays a central role in this network and is consistently and reliably activated during AM retrieval, including at the single-subject level. Several studies have investigated the relationship between hippocampal subfields and AM, albeit with mixed results (Barry et al. 2021, Bonnici et al. 2013, Miller et al. 2020, Chadwick et al. 2014, Palombo et al. 2018, Bartsch et al. 2011). For example, Bonnici (2013), Chadwick (2014), and Miller (2020) reported evidence that the CA3 region may be particularly involved in AM retrieval while Bartsch (2011) speculated a key role of the CA1 region. Furthermore, Palombo (2018) and Barry (2021) found support for a central position of the subiculum and pre/parasubiculum in AM retrieval. However, these mixed observations may result from methodological differences across these studies which used different subfield segmentation protocols (e.g., Bonnici et al 2013 did not segment the pre/parasubiculum), different imaging modalities/analysis (i.e., structural versus functional MRI), and different measures of AM retrieval success (i.e., interview-based markers versus task-based fMRI, etc). Taking this into consideration, our understanding of how hippocampal subfields preferentially engage during AM retrieval remains limited.

In addition to its subfields, functional differentiation has also been observed along the longitudinal axis of the hippocampus (Poppenk *et al*. 2013, Strange *et al*. 2014, Zeidman *et al*. 2016). Interestingly, an increasing number of 3T fMRI studies have observed that a specific region in the anterior medial hippocampus is consistently engaged during AM tasks (see Zeidman et al. 2016 for review). While the majority of these studies did not have the spatial resolution to explicitly examine hippocampal subfields, activation patterns consistently align with the medial most portion of the anterior body of the hippocampus, aligning with the location of the pre/parasubiculum and distal subiculum. Indeed, the pre/parasubiculum has recently been proposed as crucial hub for scene-based cognition (Dalton and Maguire, 2017) and subsequent experimental work has provided empirical support that this specific region preferentially engages during scene-based cognition (Dalton *et al.,* 2018; Grande *et al.,* 2022). In addition, previous research has shown that this region of the anterior medial body of the hippocampus (aligning with the location of the pre/parasubiculum) was functionally connected with parts of the AM network including the vmPFC and medial/lateral parietal cortices during the vivid re-experiencing of autobiographical memories (McCormick *et al*., 2015). Despite these preliminary insights, we do not know how different portions of hippocampal subfields along their anterior-posterior axis engage during AM retrieval.

As noted above, traditional 3T fMRI sequences at a whole-brain level and a reasonable repetition time are usually limited to a voxel size of approximately 3 mm isotropic, thus prohibiting accurate functional imaging of small brain structures like hippocampal subfields in a whole-brain setting (Willems and Henke, 2021). Technical advances in high-resolution 3T fMRI have facilitated an increase in spatial resolution that can be used to capture dissociable activity of small adjacent brain structures of the medial temporal lobes, including hippocampal subfields (Bonnici, Chadwick, Lutti, *et al*., 2012; Dalton, McCormick and Maguire, 2019; Dalton, McCormick, De Luca, *et al*., 2019). However, these sequences require a reduced field-of-view for a reasonable spatiotemporal resolution, thus precluding detailed examination of hippocampal subfield interactions with the broader AM network. Recent innovations in ultra-high field 7 Tesla (7T) fMRI sequence development including 2D high parallel imaging acceleration capabilities now permit submillimeter voxel sizes at a whole-brain level while keeping temporal resolution high (Stirnberg and Stöcker, 2021; Stirnberg *et al*., 2017). In the current study, we leveraged these technological advances to conduct a fine-grained examination of functional connectivity between hippocampal subfields along their longitudinal axis and the neocortex during AM retrieval.

To date, few studies have investigated memory signals using 7T fMRI (Berron *et al*., 2016; Grande *et al*., 2019; Risius *et al*., 2013). For example, recent reduced field-of-view 7T fMRI studies examined hippocampal subfield contributions to distinguish between or combine similar experiences (Berron *et al*., 2016; Grande *et al*., 2019). Other neuroimaging studies have exploited advances of 7T fMRI to study targeted layer-specific effects of mnemonic processes (Finn *et al*., 2019; Maass *et al*., 2014; Norris and Polimeni, 2019). However, to our knowledge, no study has hitherto leveraged both increased speed and spatial resolution of 7T fMRI to examine differential hippocampal subfield – neocortical interactions during AM retrieval.

Here, we deployed ultra-high field 7T fMRI with a dedicated AM retrieval task to achieve two primary goals: 1) to investigate the differential activation of hippocampal subfields along their longitudinal axis during AM retrieval, and 2) to examine hippocampal subfield functional connectivity with neocortical brain regions associated with the AM network.

## Materials and Methods

### Participants

24 healthy young individuals (right-handed, age: 26.66 ± 4.15 years old, Males: 12, Females: 12) with no history of neurological or psychiatric disorders and normal or corrected-to-normal vision were recruited. All participants completed secondary level of education (at least 12 years of education). All participants provided oral and written informed consent in accordance with the local ethics board.

### Autobiographical Memory and Visual Imagery Assessment

In order to examine vivid, detail-rich AM retrieval, we only included participants who reported being able to recall detail-rich personal memories and construct mental images with ease. Each participant was first asked to assess their ability to recall vivid AMs on a Likert scale from 1 to 6 (1 = able to recall detail-rich memories; 6 = unable to recall any personal events). Participants were also asked to assess their ability to construct vivid mental images on a Likert scale from 1 to 6 (1 = able to create detail-rich mental images; 6 = lack of visual imagery). This procedure was adapted from (Clark and Maguire, 2020) in which the authors demonstrated that these questions effectively captured the ability to engage in AM retrieval.

### Experimental fMRI Task

The experimental task was adapted from a previous protocol by (McCormick *et al*., 2015). The experimental procedure was clarified to the participants prior to the scanning. Participants were presented with a set of 40 randomized trials consisting of an AM retrieval task and a simple mental arithmetic (MA) task. As MA task generally does not involve the activation of hippocampus or memory, it was chosen to serve as a baseline. Each trial lasted a maximum of 17 s with a jittered inter-stimulus interval (ISI) between 1 - 4 s. The AM trials consisted of word cues of various general events, e.g., birthday celebration. Once the stimulus appeared on the screen, participants were instructed to search covertly for a relevant personal event which was specific in time and place and more than one year ago and press a response button once a memory had been chosen without verbally describing it. Participants were then asked to re-experience the memory in their mind by re-living the event with as many perceptual details as possible. In comparison, the MA trials consisted of simple addition or subtraction problems, e.g., 13 + 53. After the MA problem was solved, participants were instructed to press a response button and add 3 to the solution iteratively, e.g., (13 + 53) + 3 + 3n. Following all AM trials, participants were asked to indicate with a button press whether the AM had been re-experienced in a detailed manner or whether the retrieval was faint. Following all MA trials, participants were asked to indicate with a button press whether the MA problem had been easy or difficult.

### MRI Data Acquisition

MRI data were acquired using a MAGNETOM 7T Plus ultra-high field scanner (Siemens Healthineers, Erlangen, Germany). Participants viewed the stimulus screen placed at the back end of the scanner bore through a mirror mounted between the inner 32 channel receiver head coil and the outer circular polarized transmit coil. The MRI protocol consisted of the following scans:

#### Whole-brain T1-weighted structural image

A 0.6 mm isotropic whole-brain T1-weighted multiecho MP-RAGE scan was acquired using a custom sequence optimized for scanning efficiency (Brenner *et al*., 2013) and minimal geometric distortions (van der Kouwe *et al*., 2008) (TI = 1.1 s, TR = 2.5 s, TEs = 1.84/3.55/5.26/6.92 ms, FA = 7°, TA = 7:12, readout pixel bandwidth: 970 Hz, matrix size: 428 x 364 x 256, elliptical sampling, sagittal slice orientation, CAIPIRINHA (Breuer *et al*., 2006) 1x2_z1_ parallel imaging undersampling with on-line 2D GRAPPA reconstruction, turbofactor: 218). Finally, the four echo time images were collapsed into a single high-SNR image using root-mean-squares combination.

#### Reduced hippocampus field-of-view T2-weighted structural image

For motion-robust hippocampal subfield-segmentation, three rapid, 0.4 mm x 0.4 mm x 1.0 mm T2-weighted, slice-selective TSE scans were acquired consecutively on a reduced hippocampus field-of-view (TE = 76 ms, TR = 8090 ms, FA = 60° using Hyperecho (Hennig and Scheffler, 2001), TA = 2:59, readout pixel bandwidth: 150 Hz, matrix size: 512 x 512, 55 oblique-coronal slices of 1 mm thickness orthogonal to the long hippocampus axis, 3-fold parallel imaging undersampling with online 1D GRAPPA reconstruction, turbofactor: 9). The RF transmit power reference voltage was varied across the scans (200V, 240V, 280V) such that the nominal refocusing flip angles of the protocol were approximately reached in all brain regions in at least one of the scans. Finally, the three images from each participant were coregistered, denoised following the Rician noise estimation (Coupé *et al*., 2010) and averaged.

#### Rapid whole-brain submillimeter fMRI

A custom interleaved multishot 3D echo planar imaging (EPI) sequence was used (Stirnberg and Stöcker, 2021) with the following paramaters: TE = 21.6 ms, TR_vol_ = 3.4 s, FA = 15°, 6/8 partial Fourier sampling along the primary phase encode direction, oblique-axial slice orientation along the anterior-posterior commissure line, readout pixel bandwidth: 1136 Hz, matrix size: 220 x 220 x 140. To obtain both a high nominal spatial resolution of 0.9 mm isotropic at TR_vol_ = 3.4 s whilst imaging the whole brain at sufficient signal-to-noise-ratio (SNR) with a BOLD-optimal TE, several unique sequence features were combined for this work at 7T. A: skipped-CAIPI 3.1x7_z2_ sampling (SNR-optimized 7-fold CAIPIRINHA undersampling combined with interleaved 3-shot segmentation) (Stirnberg and Stöcker, 2021) with on-line 2D GRAPPA reconstruction. B: one externally acquired phase correction scan per volume instead of typically integrated phase correction scans per shot (Stirnberg and Stöcker, 2021). C: variable echo train lengths, skipping only the latest EPI echoes outside of a semi-elliptical k-space mask that defines 0.9 mm isotropic voxel resolution (Stirnberg *et al*., 2017). D: Rapid slab-selective binomial-121 water excitation instead of time-consuming fat saturation (Stirnberg *et al*., 2016). A 3-minute fMRI practice-run was performed before the two main functional sessions, which lasted approximately 15 min each. The MRI session was concluded by a standard 3 mm isotropic two-echo gradient-echo field-mapping scan acquired within 35 s. A maximum number of 264 imaging volumes were acquired from each functional session. The first 5 images consisting of the waiting period of 17 s prior to the beginning of the first trial were excluded to rule out non-steady-state signals.

### MRI Data Processing

#### Segmentation of Hippocampal Subfields

Manual segmentation of hippocampal subfields was performed on the averaged and denoised native space T2-weighted structural scans according to the protocol described by Dalton and colleagues (Dalton *et al*., 2017). ROI masks were created for six hippocampal subfields including DG/CA4, CA3/2, CA1, subiculum, pre/parasubiculum, and uncus using the software application ITK-SNAP 3.8 (Yushkevich *et al*., 2006). We excluded the uncus from our analyses because this region contains a mix of different hippocampal subfields (Ding and Van Hoesen, 2015) that are difficult to differentiate on structural MRI scans (Dalton *et al*., 2017) even with the high-resolution achieved in the current study. To assess intra- and inter-rater reliability, five hippocampi were segmented by two independent raters (P.L. and M.A.D) and again six months after initial segmentation. The inter-rater reliability as measured by the DICE overlap metric (Dice, 1945) were in accordance with those reported in the existent literature (Bonnici, Chadwick, Kumaran, *et al*., 2012; Palombo *et al*., 2013): DG/CA4 = 0.87 (aim 0.86-0.80), CA3/2 = 0.73 (aim of 0.74-0.67), CA1 = 0.80 (aim of 0.81-0.67), subiculum = 0.80 (aim of 0.79-0.57), and pre/parasubiculum = 0.64 (aim of 0.67-0.57). Intra-rater reliability was measured 8 months apart and also showed high concordance between segmentations at the two different time points (0.92 for DG/CA4, 0.79 for CA3/2, 0.84 for CA1, 0.84 for subiculum, and 0.86 for pre/parasubiculum).

#### Segmentation of Hippocampal Subfields in Anterior, Anterior Body, Posterior Body, and Tail Portions

Each hippocampal subfield ROI mask was divided into four portions along the longitudinal axis of the hippocampus (head, anterior body, posterior body, and tail) using the software application ITK-SNAP 3.8, according to the protocol described by Dalton and colleagues (2017; 2019). The head masks encompassed the first slice on which the hippocampus was visible up to the slice preceding the first slice of the DG. The mean number of slices in the head mask was 7.67 (SD = 2.01). The remaining slices, beginning with the first slice of the DG until the last slice of the hippocampus, were summed and divided into 3 parts to create equivalent slices in the anterior body, posterior body, and tail masks, resulting in a mean of 11.04 (SD = 1.01), 10.73 (SD = 0.98), and 10.54 (SD = 0.94) slices respectively. The average total number of slices covering the hippocampus was 39.98 (SD = 2.57).

#### Functional MRI preprocessing

To address both research aims, two steps of MRI preprocessing were performed using SPM12 (Statistical Parametric Mapping 12) software package (www.fil.ion.ucl.ac.uk/spm/) on MATLAB v17a (MathWorks) computing platform (https://matlab.mathworks.com/). For both steps, the anatomical and the functional scans obtained from each subject were reoriented to be in line with the anterior-posterior commissure axis. The field maps, including the phase and magnitude images, were used to calculate the voxel displacement maps (VDM) to geometrically correct the distorted EPI images. The VDMs were then applied to the functional scans during realignment and unwarping. The averaged anatomical scans (and hippocampal subfield masks) were co-registered with the functional scans. After motion correction and co-registration, the preprocessing pipelines for the two research aims diverged: For the first aim (to examine differential activation of hippocampal subfields), fMRI data were kept in native space to allow maximum spatial precision. Only a sparse Gaussian smoothing kernel of 1 mm full-width half-maximum (FWHM) to reduce excess noise (Yoo *et al*. 2018) and a temporal high-pass filter of 128 s was applied to the function data. Then, one-sample T-Test contrasts (T-contrasts) were calculated to test the effects of AM retrieval versus baseline (MA). For the second aim (to examine functional connectivity of hippocampal subfields to other neocortical regions), motion-corrected and co-registered fMRI data were normalized to the Montreal Neurological Institute (MNI) space and smoothed with a slightly smaller than standard 6 mm FWHM to examine activation and functional connectivity at the group level.

Additionally, in order to mitigate the artifacts produced by unsolicited movements during the functional tasks for further exclusion criteria, we screened for motion artifact outliers in the time series using the ARtifact detection Tools (ART) software package (v2015). The standard threshold of outlier detection if motion exceeds the 97th percentile of the global mean intensity (relating to under 1 mm motion) for more than 10% of the number of the scans. No excessive motion was detected, hence, no participant was excluded from the study.

### Statistical Analyses

#### Hippocampal Subfield Activation Extraction

For the native space fMRI analyses, we followed the standard GLM procedure in SPM12 with trials designated as mini blocks and covering the elaboration period fixed at the last 10 s of the stimulus time and prior to the display of the vividness rating. Since only a few trials were rated as faint/difficult, we included all trials in our analyses. Furthermore, motion correction parameters were included in the GLM as covariate of no interest. The contrasts of interest of the first level were specified as 1) AM versus Baseline and 2) MA versus Baseline. Signal intensity values were extracted for both contrasts for the five right and five left hippocampal subfield ROIs covering the entire length of the hippocampus. We then subtracted the signal intensities for MA from the signal intensities for AM for each of the ROIs for each participant. In a second step, we extracted signal intensities for AM and MA of the five right and five left hippocampal subfields for the anterior body, posterior body, and tail separately. Signal intensities were extracted for each participant in native space using MATLAB-based Response Exploration (REX) toolbox (https://www.nitrc.org/projects/rex) by applying the segmented ROI masks. Since we found no evidence of laterality effects (t = 1.04, df = 23, p = 0.3568, see Table S1), signal intensities for bilateral subfields were collapsed. Differential signal intensity values were subjected to a 1-way-RM-ANOVA with Tukey’s multiple comparison test. A significance threshold of p < 0.05 was applied. Furthermore, the temporal signal-to-noise ratios (tSNR) across the fMRI time series along the longitudinal axis of the hippocampal subfields were examined and the 1-way-RM-ANOVA with Dunnett’s multiple comparison tests were applied (see Figure S2).

#### Group analyses of whole-brain activation and hippocampal subfield functional connectivity

First, to assess differences between AM and MA, a multivariate mean-centered partial least squares (PLS) group analysis was performed. Detailed descriptions of PLS can be found elsewhere (Krishnan *et al*., 2011; McIntosh and Lobaugh, 2004). In brief, PLS uses singular value decomposition to extract ranked latent variables (LVs) from the covariance matrix of brain activity and conditions in a data driven manner. These LVs express patterns of brain activity associated with each condition. Statistical significance of the LVs was assessed using permutation testing. In this procedure, each participant’s data was randomly reassigned (without replacement) to different experimental conditions, and a null distribution was derived from 500 permutated solutions. We considered LVs as significant at p < 0.05. Furthermore, we assessed the reliability of each voxel that contributed to a specific LV’s activity pattern using a bootstrapped estimation of the standard error (bootstrap ratio, BSR). For each bootstrapped solution (100 in total), participants were sampled randomly with replacement and a new analysis was performed. In the current study, we considered clusters of 50 or more voxels with BSRs greater than 3 (approximately equal to a p < 0.001) to represent reliable patterns of activation. Of note, PLS uses two re-sampling techniques that (1) scramble the data of each participant’s conditions so that small but reliable differences between true experimental conditions can be detected, and (2) exclude whole datasets of participants, so that outliers who may drive significant effects can be detected.

To assess functional connectivity between hippocampal subfields and the rest of the brain, seed PLS, an extension of the mean-centered PLS was used (McIntosh and Lobaugh, 2004). Seed PLS examines the relationship between a target region (seed region) and signal intensities in all other brain voxels as a function of the experimental conditions (Krishnan et al. 2011). The main difference to the mean-centered PLS is that, in seed PLS, the covariance matrix used in the single value decomposition stems from correlation values between the seed region and all other voxels for each experimental condition. Thus, seed PLS offered us to examine the multivoxel patterns which correlate with fMRI signal extracted from individual hippocampal subfields. Signal intensities of individual subfields (extracted in native space) were used as seeds for the group analyses (performed in MNI space). We conducted three seed PLS analyses (1. anterior body, 2. posterior body, and 3. Tail) containing all five hippocampal subfields and contrasting AM vs MA trials. A significance threshold for the LV’s (500 permutations) of p < 0.05 was. After establishing whether the functional connectivity pattern differed between AM and MA across all five subfields for a specific portion of the hippocampal long-axis, we then examined this portion more closely with follow-up PLS analyses. In these post hoc analyses, functional connectivity of each of the five hippocampal subfields were assessed separately and a Bonferroni multiple comparison correction was applied so that we considered statistical significance for the LV’s at p < 0.01 (see for a similar approach (McCormick *et al*., 2021)). Boot strap ratios (100 bootstraps) of < 3 and > 3 (corresponding approximately to a p < 0.01) were considered significant.

## Results

### Behavioural Results

All 24 participants reported being able to recall detail-rich autobiographical memories (average of 1.92 ± 0.67, 1 = able to recall detail-rich memories; 6 = unable to recall any personal events) and construct vivid mental images (average of 1.83 ± 0.69, 1 = able to create detail-rich mental images; 6 = lack of visual imagery). During scanning, the participants spent 3.53 s (± 0.98) on average to select a memory and around 3.66 s (± 0.97) to solve the MA problem. Whereas some trials were excluded from the analyses due to missing button presses, no significant difference was found between the speed of AM retrieval and MA solving (t = 0.8601, p = 0.3958). Participants indicated in 35.67 (± 4.67) trials out of 40 trials that their memories were vivid (t = 19.36, df = 23, p < 0.0001) and 28.13 (± 6.15) trials out of 40 MA problems were reported as easy (t = 8.553, df = 23, p < 0.0001).

### Hippocampal Subfield Activation During AM

Greater bilateral hippocampal activation during AM retrieval than MA solving was found in all participants (see Figure 1). Furthermore, all hippocampal subfields showed greater activation during AM retrieval than solving a MA problem (DG/CA4: t = 6.140, df = 23, p < 0.001, CA2/3: t = 5.217, df = 23, p < 0.001, CA1: t = 3.768, df = 23, p = 0.001, subiculum: t = 3.068, df = 23, p = 0.005, pre/parasubiculum: t = 4.637, df = 23, p < 0.001, Bonferroni corrected). Further analyses revealed differences in hippocampal subfield activity associated with AM retrieval (F = 5.887, df = 5, p = 0.017, Figure 1 and Table S1 for an overview of % signal changes). In line with our hypothesis (Figure 1E), we found that activation during AM was much stronger in the pre/parasubiculum compared with CA1 (df = 23, p = 0.001), the subiculum (df = 23, p = 0.001), CA2/3 (df = 23, p = 0.049), and at a non-significant trend level in DG/CA4 (df = 23, p = 0.075). There were no other significant differences between subfield activation associated with AM retrieval.

**Figure 1.**
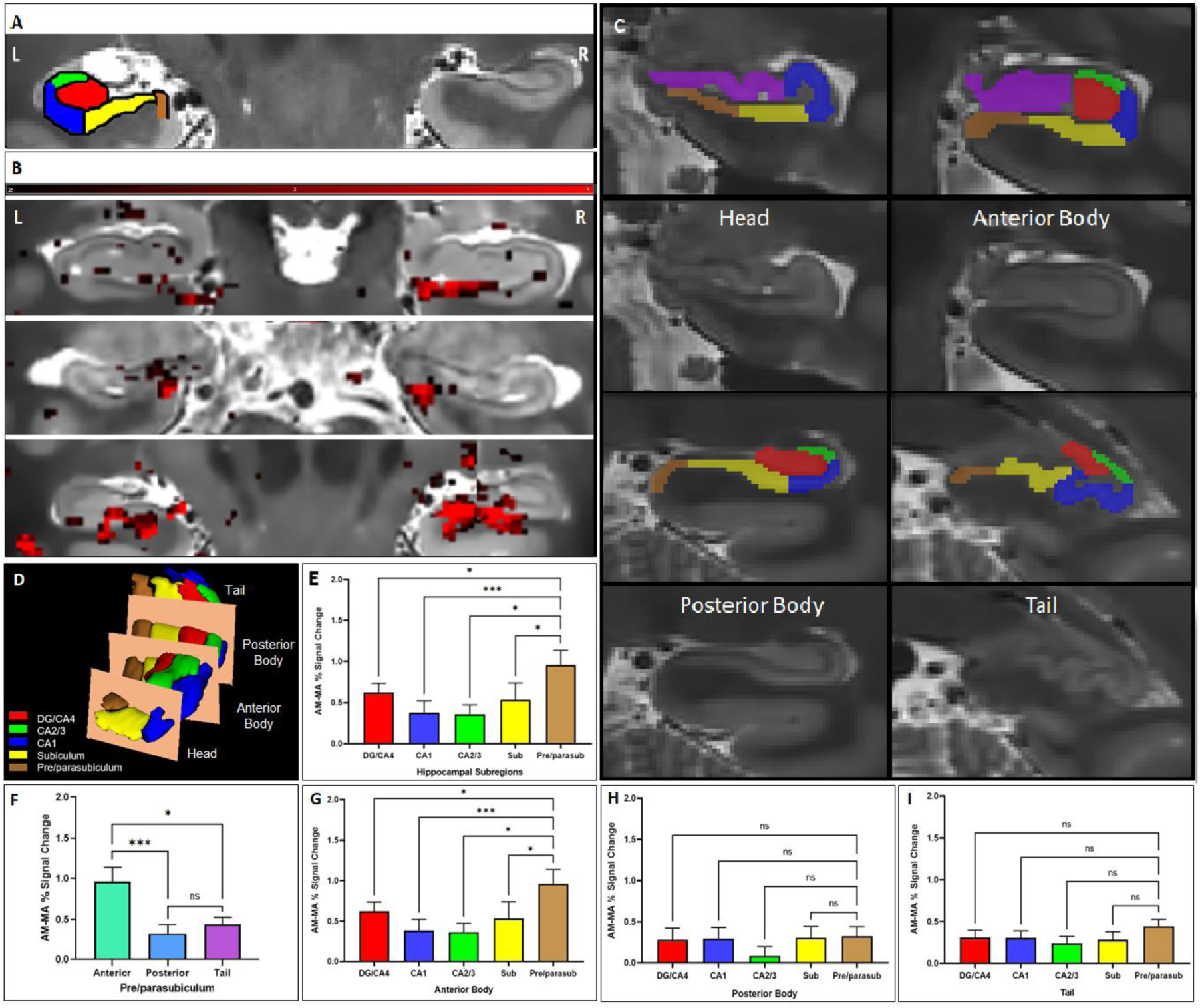
(Color). Differential hippocampal subfield engagement during AM retrieval. A. Overlaid segmentation of labelled hippocampal subfields, including the DG, CA1-4, subiculum, and pre/parasubiculum on high-resolution structural T2-weighted scan. B. Examples of AM versus MA activation along the longitudinal axis of the hippocampus (shown in red) from three participants (Y coordinates from upper to lower panels of 20, 17, and 27, beginning from rostral to caudal of 55 slices, respectively). C. Examples of manual hippocampal subfields segmentation for signals extraction. The subfields along the longitudinal axis, are divided into 4 portions of head, anterior body, posterior body, and tail (Y coordinates of 16, 21, 38, and 46, respectively). D. Hippocampus subfields along the longitudinal axis. E. The comparison between the % signal change during AM and MA in hippocampal subfields (DG/CA4, CA1, CA2/3, subiculum, and pre/parasubiculum). The pre/parasubiculum showed stronger differentiation between AM and MA than most other subfields. F. This effect was driven by the anterior body part of the hippocampus. G. The anterior body of the pre/parasubiculum shows greater differentiation between AM and MA than all other subfields, whereas no significant difference was found in the posterior body (H) nor the tail (I). *** p < 0.001, ** p < 0.01, and * p < 0.05, and + p < 0.1 (non-significant trend level).

### Hippocampal Subfield Activation along its longitudinal axis during AM

Next, we assessed differential subfield engagement of the anterior body, posterior body, and tail separately. Strikingly, the RM-ANOVA found a main effect of subfield activation levels in the anterior body portion of the hippocampus (F = 4.440, df = 4, p = 0.024, see Figure 1G) but not in the posterior body (F = 1.650, df = 4, p = 0.1895) or tail (F = 1.157, df = 4, p = 0.3286) of the hippocampus (Figure 1H and 1I, respectively). The head portion was omitted from these analyses due to 1) significantly less number of slices and voxel count compared to the other portions and 2) not all subfields being present in this portion of the hippocampus (e.g., the DG). No signal intensity was detected in the head portion in one participant, possibly due to signal drop out. Post hoc analyses revealed that, in the anterior body portion of the hippocampus, activation relating to AM versus MA was much stronger in the pre/parasubiculum than in CA1 (df = 23, p < 0.001), CA2/3 (df = 23, p = 0.019), subiculum (df = 23, p = 0.034), and DG/CA4 (df = 23, p = 0.034). There were no other significant differences between anterior body subfields in activation associated with AM retrieval. Further comparison (F(2, 23) = 9.878, p < 0.001) revealed that the activation difference (AM vs MA) of the pre/parasubiculum in the anterior body was stronger than the pre/parasubiculum in the posterior body (df = 23, p < 0.001) and the tail (df = 23, p = 0.020).

### Hippocampal Subfield - Neocortical Interactions During AM

On a whole-brain, whole-group basis, we found greater activation during AM retrieval than MA in all regions typically associated with AM retrieval, including bilateral hippocampal activation as well as vmPFC and medial/lateral parietal activation (LV1, p < 0.0001, see Figure 2 and Table S3 for peak MNI coordinates).

**Figure 2.**
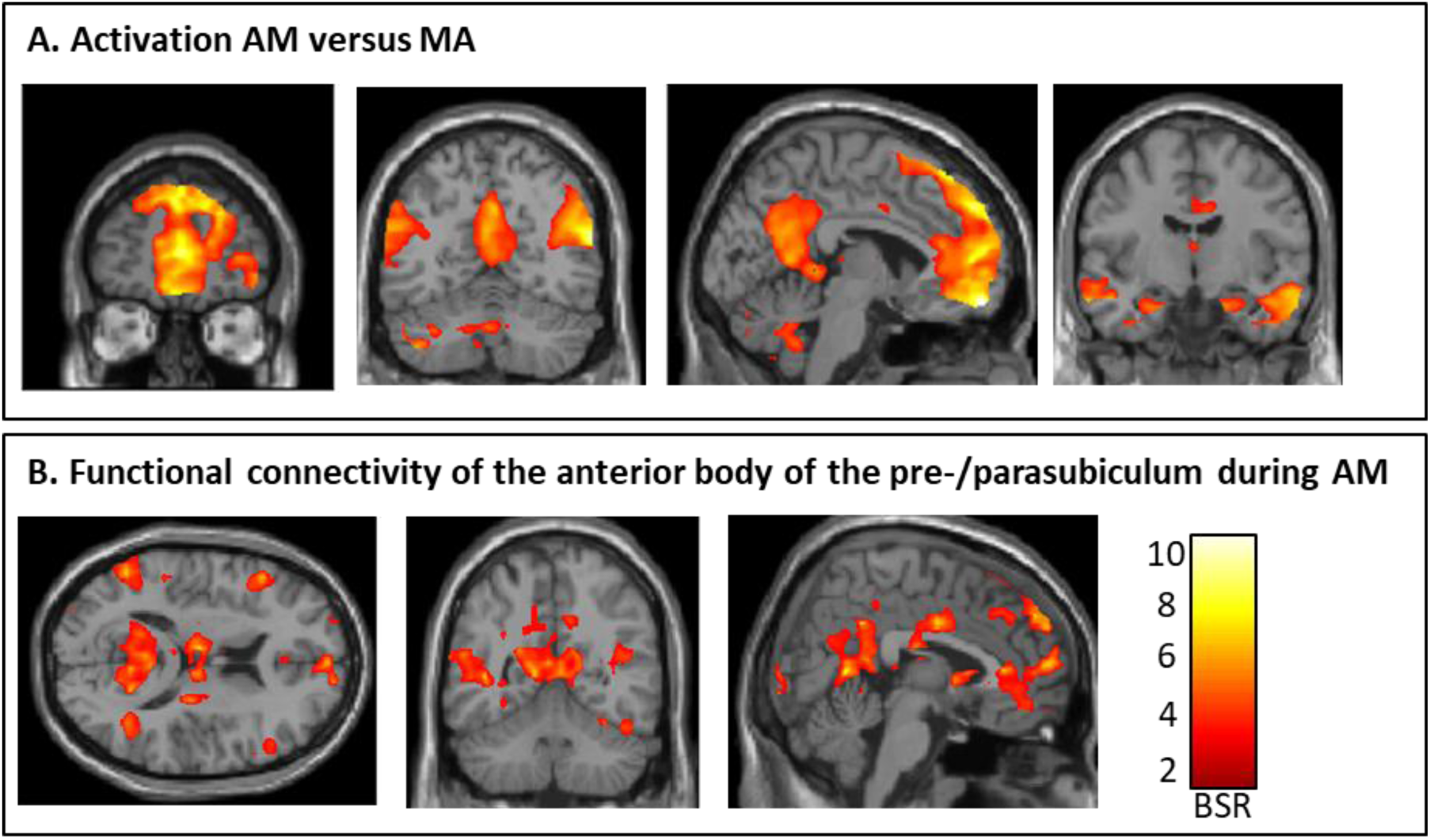
Hippocampal subfield functional connectivity during AM. A. Greater activation for AM versus MA is shown in bootstraps ratios (BSR). All regions typically associated with AM show greater activation of bilateral hippocampi, vmPFC and medial/lateral parietal cortex for AM. The whole-brain fMRI activation is overlaid on a standard T1-weighted MRI image. B. Greater functional connectivity of the anterior body of the pre/parasubiculum during AM than MA is shown. All regions typically associated with AM retrieval show strong functional connectivity, including vmPFC and medial/lateral parietal cortices.

Next, we evaluated patterns of functional connectivity along the long-axis of the hippocampus. We found that only functional connectivity patterns of the anterior body of the hippocampus differed between AM and MA (anterior body: LV 1, p = 0.027, posterior body: LV 1, p = 0.51, tail: LV 1, p = 0.64). In Bonferroni corrected post hoc analyses (thus applying a threshold of p < 0.01), we found that only the functional connectivity of the pre/parasubiculum differed between both conditions (pre/parasubiculum: p < 0.001), whereas the other subfields did not (DG/CA4: p = 0.53, CA2/3: p = 0.16, CA1: p = 0.36, subiculum: p = 0.28). Figure 3 illustrates the brain pattern associated with functional connectivity of the anterior body of the pre/parasubiculum during AM, including all canonical regions typically associated with AM, including the vmPFC and medial/lateral parietal cortices (see Table S4 for peak MNI coordinates). In addition to these areas that are traditionally associated with the AM network, the anterior body of the pre/parasubiculum also showed significant functional connectivity with posterior visual-perceptual cortices (i.e., right lingual gyrus and bilateral fusiform gyrus) during AM than MA.

## Discussion

By exploiting a novel whole-brain 7T fMRI sequence with submillimeter voxelsize, this study provides two novel insights into hippocampal memory processes. First, we found evidence that the anterior body of the pre/parasubiculum was significantly more engaged during the vivid re-experience of AMs than other hippocampal subfields. Second, during AM, only the anterior body of the pre/parasubiculum showed stronger functional connectivity to neocortical brain regions typically engaged during AM retrieval over and above other hippocampal subfields. We discuss these findings in turn. We observed that all hippocampal subfields differentiated between AM and MA with the pre/parasubiculum differentiating between cognitive tasks to a greater extent than the neighboring subiculum and CA fields. While our AM task was not designed to specifically target different cognitive states (i.e., scene-specific content), our finding indicates that overall AM retrieval preferentially engages the pre/parasubiculum. Examining our study design closer, we only included participants who reported to be able to retrieve visually detail-rich AMs easily and we focused our analyses on the last 8 secs of the AM trials. This served to highlight the period in which participants were most likely engaged in re-experiencing visual-perceptual imagery. One explanation for our findings, therefore is that the pre/parasubiculum is preferentially engaged in tasks which rely on vivid scene-based cognition (Dalton and Maguire, 2017). For example, previous work suggests that the pre/parasubiculum is more strongly activated during scene-imagery than object-imagery (Dalton *et al*., 2018; Zeidman *et al*., 2015), and scene/object discrimination (Hodgetts *et al*., 2017). Furthermore, we know from rodents and nonhuman primates that the pre/parasubiculum contains an abundance of head, grid, and border cells (Boccara *et al*., 2010; Lever *et al*., 2009; Robertson *et al*., 1999), which has recently been extended to human goal direction cells (Shine *et al*., 2019). Arguably, AM retrieval relies heavily on scene-based cognition since AMs tend to unfold on a visuo-spatial stage (Greenberg and Knowlton, 2014). In fact, when participants are asked to imagine personal events, they tend to place the event onto a spatial stage (Robin *et al*., 2018), indicating that visuo-spatial imagery plays a fundamental role in episodic memory retrieval. Further strengthening the tight link between the ability to recall AMs and visual imagery, people with little or no ability to experience visual imagery, commonly referred to as aphantasics, also tend to have difficulties recalling AM (Dawes *et al*., 2020; Milton *et al*., 2021; Zeman *et al*., 2015, 2020). Interestingly, aphantasics not only have difficulties to recall visual-perceptual details to their AMs, but they seem to have a worse ability to retrieve personal memories in general (Dawes *et al*., 2020) and show poorer verbal and non-verbal memory function (Monzel *et al*., 2021). In fact, a recent phenomenon called severe deficient autobiographical memory or SDAM (Palombo, Sheldon, *et al*., 2018; Watkins, 2018) has also been associated with aphantasia (Pearson, 2019). This line of thought meshes well with the scene construction theory positing that a dominant function of the hippocampus is to construct spatially coherent internal models of the environment (Dalton and Maguire, 2017; Maguire and Mullally, 2013; McCormick, Ciaramelli, *et al*., 2018; Zeidman and Maguire, 2016), and the pre/parasubiculum may be of special significance to this process (Dalton and Maguire, 2017; Dalton *et al*., 2018). In line with the scene construction theory, patients with hippocampal damage have been shown to use less scene-based cognition in their mind-wandering episodes (McCormick, Rosenthal, *et al*., 2018), moral decision-making (McCormick *et al*., 2016), and scene-based judgements (McCormick *et al*., 2017). Therefore, our results point towards a potential role of the pre/parasubiculum in tasks relying heavily on vivid visuo-spatial imagery, such as vivid AM retrieval.

Having highlighted the role of the pre/parasubiculum in vivid AM retrieval, we further found that all other hippocampal subfields also showed stronger activation during AM retrieval than MA. This result is not surprising since the contrast between AM and MA is loose and differences between the two cognitive states may not be specific to any one process. Especially, since AM retrieval is a complex cognitive task with a magnitude of different operations (such as detail integration, and discrimination), it is likely that the other hippocampal subfields contribute to different processes which we could not dissociate in the current study. In fact, recent investigations show different contributions of hippocampal subfields to mnemonic processes. Although there is somewhat mixed evidence in the current literature, CA fields have been implicated in the integration or associations of memory details, such as external and internal (Chadwick *et al*., 2014; Grande *et al*., 2019; Miller *et al*., 2020; Newman and Hasselmo, 2014), whereas the DG/CA4 region (Baker *et al*., 2016; Berron *et al*., 2016; Newman and Hasselmo, 2014; van Dijk and Fenton, 2018) may support the separation or discrimination of mnemonic information. The aim of the current study was to evaluate a submillimeter 7T fMRI sequence during a robust, reliable, and established AM paradigm. Future studies will now have to experimentally target specific subfield functions, such as examining AMs with and without visual imagery. Our newly developed 7T fMRI sequence will allow the innovative investigation of differential hippocampal subfield contributions to cognition.

The second major goal of the current study was to examine functional connectivity of hippocampal subfields to the neocortex during AM retrieval. We found that the anterior body of the pre/parasubiculum, over and above other subfields, has strong functional connectivity to neocortical regions known to support AM retrieval. In addition, this effect was specific to the anterior body and not evident in the posterior body or tail portion of the hippocampus. This finding adds new detail to several lines of research. For example, both 3T and 7T fMRI resting state studies have identified the subiculum (in which the pre/parasubiculum was included) as a functional connectivity hub correlating with activity in the default mode network which has overlapping brain regions to the AM network (Ezama *et al*., 2021; Shah *et al*., 2018). In addition, previous task-based 3T fMRI revealed that hippocampal functional connectivity during AM of a seed region in the vicinity of the pre/parasubiculum (McCormick *et al*., 2015) was strongly connected to a brain-wide network comprising the vmPFC and medial/lateral parietal cortices, as well as visual-perceptual regions of the occipital cortex. Furthermore, the same region of the anterior medial hippocampus was more strongly connected to frontal and parietal cortices during scene construction than object construction (Zeidman *et al*., 2015). Additionally, from a neuroanatomical perspective, the pre/parasubiculum is a primary target of the parieto-medial temporal visual pathway carrying information about visuospatial representations of the environment (Dalton and Maguire, 2017). The parieto-medial temporal pathway directly links the pre/parasubiculum with the inferior parietal lobule, posterior cingulate cortex, retrosplenial cortex, and parahippocampal gyrus (Ding and Van Hoesen, 2015; Ding, 2013; Kravitz *et al*., 2011). Each of these regions have been heavily associated with visuospatial cognition (Auger and Floresco, 2014; Epstein *et al*., 2007) and AM (Svoboda *et al*., 2006) and connect directly with the pre/parasubiculum, giving it privileged access to visuo-spatial information. While most of this evidence stems from anatomical connectivity studies in rodent and nonhuman primates, a recent diffusion weighted imaging study supports this framework by showing, for the first time in the human brain, that circumscribed regions along the anterior-posterior axis of the pre/parasubiculum, including a specific portion in the anterior body of the hippocampus, have dense patterns of anatomical connectivity with distributed cortical brain areas implicated in AM (Dalton *et al*., 2022). The anterior portion was shown to exhibit greater connectivity with temporal, medial parietal, and occipital regions. The posterior hippocampus, more intense in the tail, was partially found to be connected with medial parietal and occipital cortices. Our results dovetail nicely with this collection of structural and functional data and provide new evidence that the pre/parasubiculum in the anterior body of the hippocampus may be an important hippocampal hub for scene-based cognition.

In summary, here we utilized a novel submillimeter 7T fMRI sequence which enabled us to examine functional connectivity between hippocampal subfields and neocortical regions during vivid AM retrieval. We enhanced our knowledge of hippocampal subfield contributions to cognition by showing that the anterior body of the pre/parasubiculum was more engaged during AM than other neighboring hippocampal subfields and that this part of the hippocampus was strongly functionally connected to regions typically recruited during AM. In context of the broader literature, these observations correspond well with multiple lines of evidence that suggest a specific portion of the pre/parasubiculum in the anterior body of the hippocampus may be a central component of networks underpinning AM retrieval and a crucial area underpinning our ability to engage in vivid AM and more broadly, scene-based cognition.

## Data/code availability

The data are available upon request by contacting the Lead Contact, Cornelia McCormick.

## Author contribution

PL: Data curation, formal analysis, original draft writing. MAD: Methodology, visualization, writing, reviewing, editing. RS: Methodolody, TS: Methodology, ASp: reviewing, editing, ASc: reviewing, editing, CM: conceptualization, methodology, data curation, visualization, supervision, writing, reviewing, editing.

## Acknowledgments

CMC and PL are supported by a BONFOR fellowship (2019-2-06, 2021-2A-01) and the German Research Foundation (MC-244/3-1). We thank Sascha Brunheim, Anke Rühling, and Jennifer Schlee for their technical support during scanning and all participants for their time.

The authors have no competing interests to declare.

## Supplementary Materials

**Figure S1.**
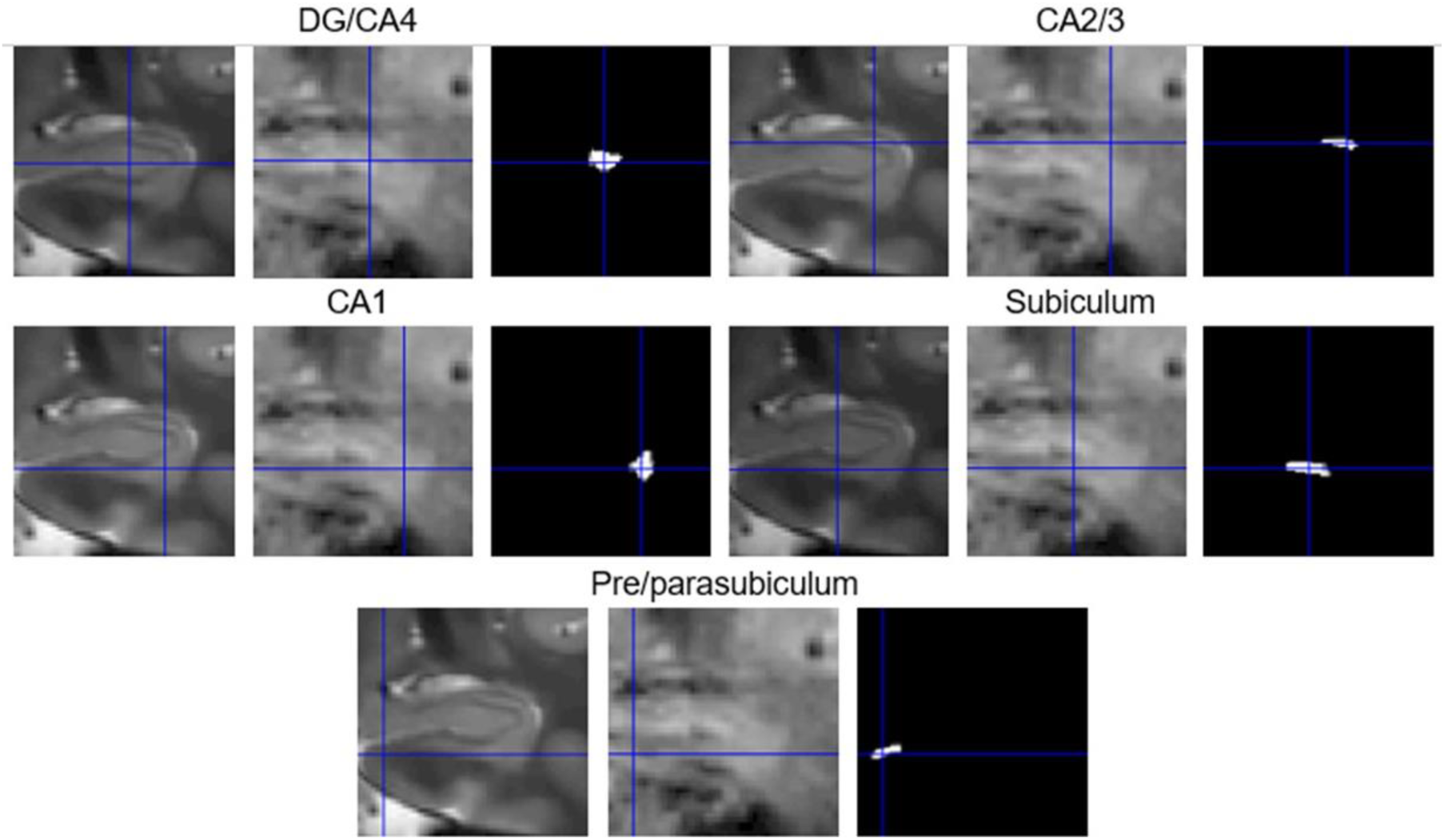
Examples of co-registration between structural and functional scans of five hippocampal subfields. High-resolution structural T2-weighted scans (left) are shown in alignment with their corresponding functional EPI scans (centre) and the ROI masks (right) in coronal view in the anterior body portion of the hippocampus. The mask of each subfield was constructed using ITK-Snap.

**Figure S2.**
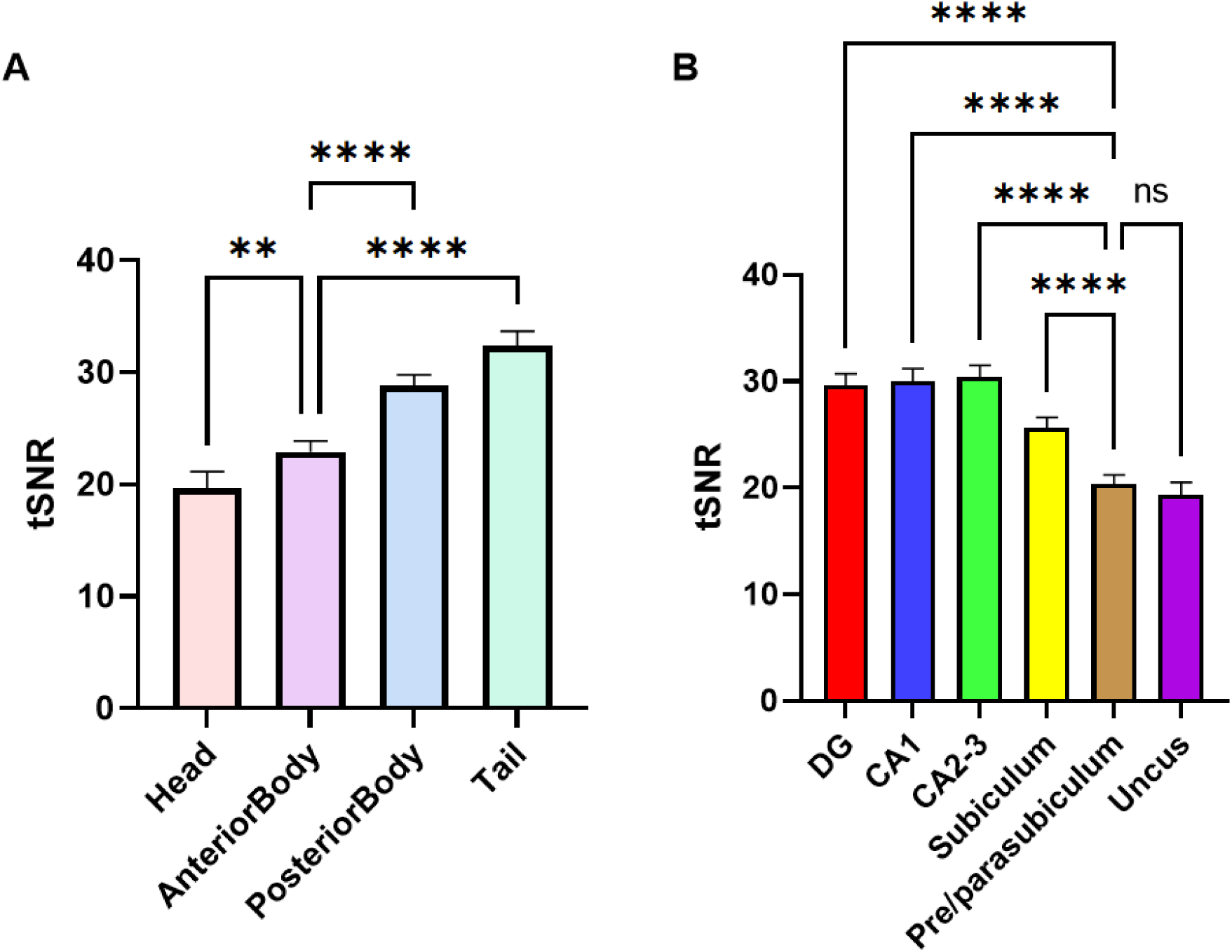
Comparison of the temporal signal-to-noise ratio (tSNR) across the fMRI time series along the longitudinal axis of the hippocampal subfields. A. The tSNR values ranged from 8.44 to 44.82 and multiple comparisons between the four hippocampal portions showed that the tSNR along the more anterior portions were significantly lower than the more posterior portions (F (1.722, 39.04) = 65.84, p < 0.001). Additionally, the pre/parasubiculum, as the most medial subfield, displayed lower tSNR than the other subfields (F (2.551, 58.68) = 129.2, p < 0.001). A possible explanation for this occurrence is the typical low SNR in the medial and lower regions of the brain including the hippocampus and its subfields. However, low tSNR values usually make it more difficult to detect significant differences between experimental conditions. Although we cannot rule out effects of tSNR differences on our results, low tSNR of the pre/parasubisulum was present during both experimental conditions. Therefore, despite low tSNR, we still find significant differences in this hippocampal subfield and not in the adjacent subiculum or posterior body of the pre/parasubiculum.

**Table S1.** Percentage of signal change during AM versus MA tasks.

|  | *Left* |  | *Right* |  | *T-Test* |
| --- | --- | --- | --- | --- | --- |
|  | **M** | **SD** | **M** | **SD** | **P-value** |
| ***Anterior*** | 0.8046 | 1.4062 | 0.7908 | 1.1998 | 0.8638 |
| ***Anterior Body*** | 0.9250 | 1.3697 | 0.6253 | 0.5293 | 0.1579 |
| ***Posterior Body*** | 0.4770 | 0.7195 | 0.3527 | 0.4562 | 0.1030 |
| ***Tail*** | 0.2187 | 0.3552 | 0.2164 | 0.5201 | 0.9774 |
| ***DG/CA4*** | 0.5416 | 0.5722 | 0.4036 | 0.3184 | 0.1258 |
| ***CA2/3*** | 0.4184 | 0.4246 | 0.1789 | 0.4936 | 0.1230 |
| ***CA1*** | 0.3741 | 0.6368 | 0.3381 | 0.4099 | 0.6119 |
| ***Subiculum*** | 0.4666 | 0.8840 | 0.4288 | 0.7922 | 0.4835 |
| ***Pre/parasubiculum*** | 0.7971 | 0.829 | 0.7091 | 0.7773 | 0.4313 |

The extracted percentages of change in activation intensity between the AM and MA in four hippocampal portions along the long axis (anterior, anterior body, posterior body, and tail) and five hippocampal subfields (DG/CA4, CA1, CA2/3, subiculum, and pre/parasubiculum), both left and right, were averaged. M= Mean, SD = Standard deviation, df = 23

**Table S2.** Percentage of signal change during AM versus MA tasks in the anterior body.

|  | *Left* |  | *Right* |  | *T-Test* |
| --- | --- | --- | --- | --- | --- |
|  | M | SD | M | SD | P-value |
| ***DG/CA4*** | 0.7749 | 0.8829 | 0.4734 | 0.5514 | 0.1481 |
| ***CA2/3*** | 0.5806 | 0.6838 | 0.1647 | 1.029 | 0.1564 |
| ***CA1*** | 0.4883 | 1.035 | 0.2945 | 0.4198 | 0.2181 |
| ***Subiculum*** | 0.6597 | 1.228 | 0.3989 | 0.8272 | 0.0348* |
| ***Pre/parasubiculum*** | 0.9746 | 0.9099 | 0.9117 | 0.9740 | 0.6290 |

The extracted percentages of change in activation intensity between the AM and MA exclusively in the five hippocampal subfields (DG/CA4, CA1, CA2/3, subiculum, and pre/parasubiculum) of the anterior body along the longitudinal axis, both left and right, were averaged. M= Mean, SD = Standard deviation, df = 23, \* p < 0.05

**Table S3.**
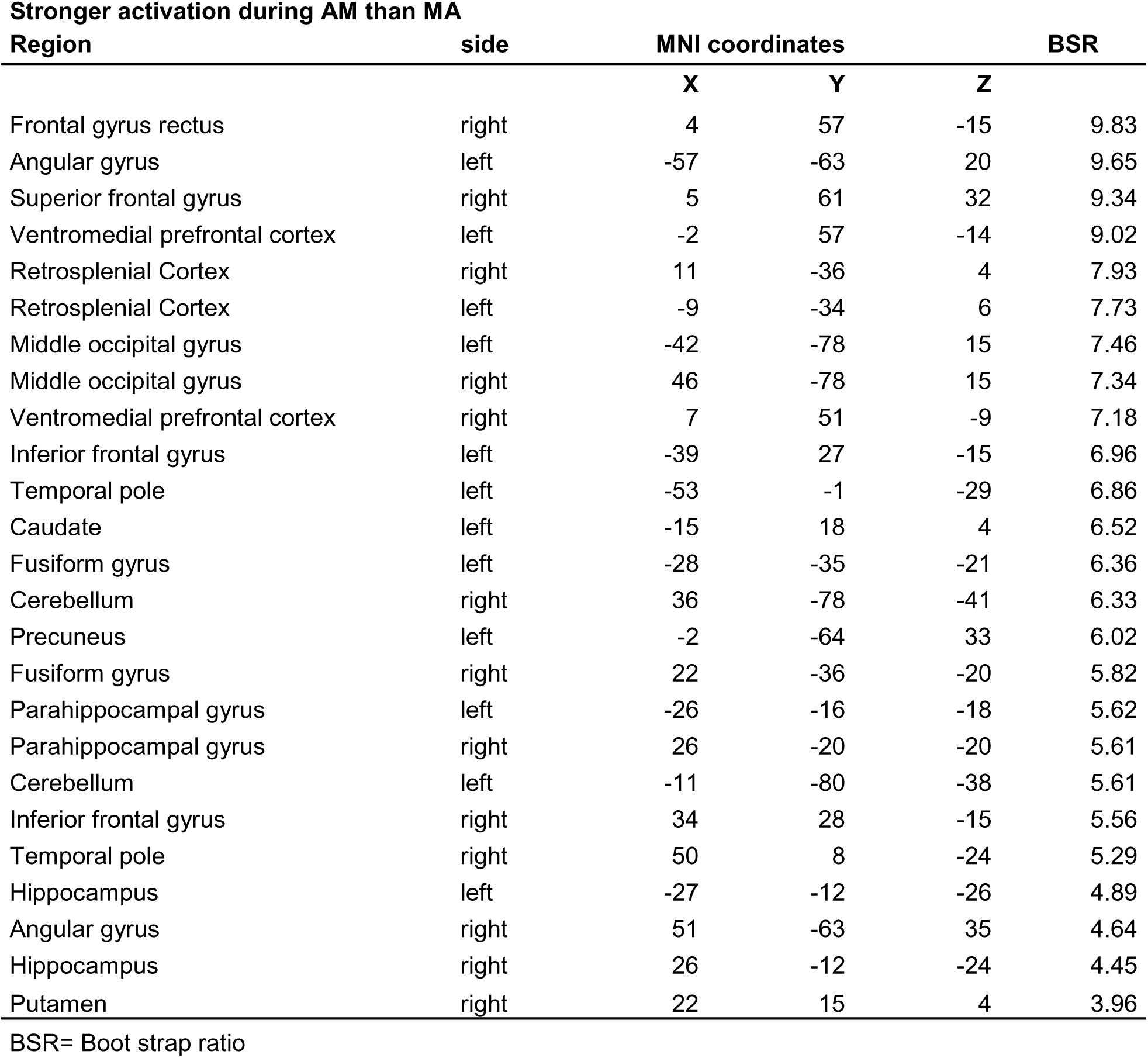
Peak coordinates of the mean-centered PLS.

| **Stronger activation during AM than MA** |  |  |  |  |  |
| --- | --- | --- | --- | --- | --- |
| **Region** | **side** | **MNI coordinates** |  |  | **BSR** |
|  |  | **X** | **Y** | **Z** |  |
| Frontal gyrus rectus | right | 4 | 57 | -15 | 9.83 |
| Angular gyrus | left | -57 | -63 | 20 | 9.65 |
| Superior frontal gyrus | right | 5 | 61 | 32 | 9.34 |
| Ventromedial prefrontal cortex | left | -2 | 57 | -14 | 9.02 |
| Retrosplenial Cortex | right | 11 | -36 | 4 | 7.93 |
| Retrosplenial Cortex | left | -9 | -34 | 6 | 7.73 |
| Middle occipital gyrus | left | -42 | -78 | 15 | 7.46 |
| Middle occipital gyrus | right | 46 | -78 | 15 | 7.34 |
| Ventromedial prefrontal cortex | right | 7 | 51 | -9 | 7.18 |
| Inferior frontal gyrus | left | -39 | 27 | -15 | 6.96 |
| Temporal pole | left | -53 | -1 | -29 | 6.86 |
| Caudate | left | -15 | 18 | 4 | 6.52 |
| Fusiform gyrus | left | -28 | -35 | -21 | 6.36 |
| Cerebellum | right | 36 | -78 | -41 | 6.33 |
| Precuneus | left | -2 | -64 | 33 | 6.02 |
| Fusiform gyrus | right | 22 | -36 | -20 | 5.82 |
| Parahippocampal gyrus | left | -26 | -16 | -18 | 5.62 |
| Parahippocampal gyrus | right | 26 | -20 | -20 | 5.61 |
| Cerebellum | left | -11 | -80 | -38 | 5.61 |
| Inferior frontal gyrus | right | 34 | 28 | -15 | 5.56 |
| Temporal pole | right | 50 | 8 | -24 | 5.29 |
| Hippocampus | left | -27 | -12 | -26 | 4.89 |
| Angular gyrus | right | 51 | -63 | 35 | 4.64 |
| Hippocampus | right | 26 | -12 | -24 | 4.45 |
| Putamen | right | 22 | 15 | 4 | 3.96 |

BSR= Boot strap ratio

**Table S4.**
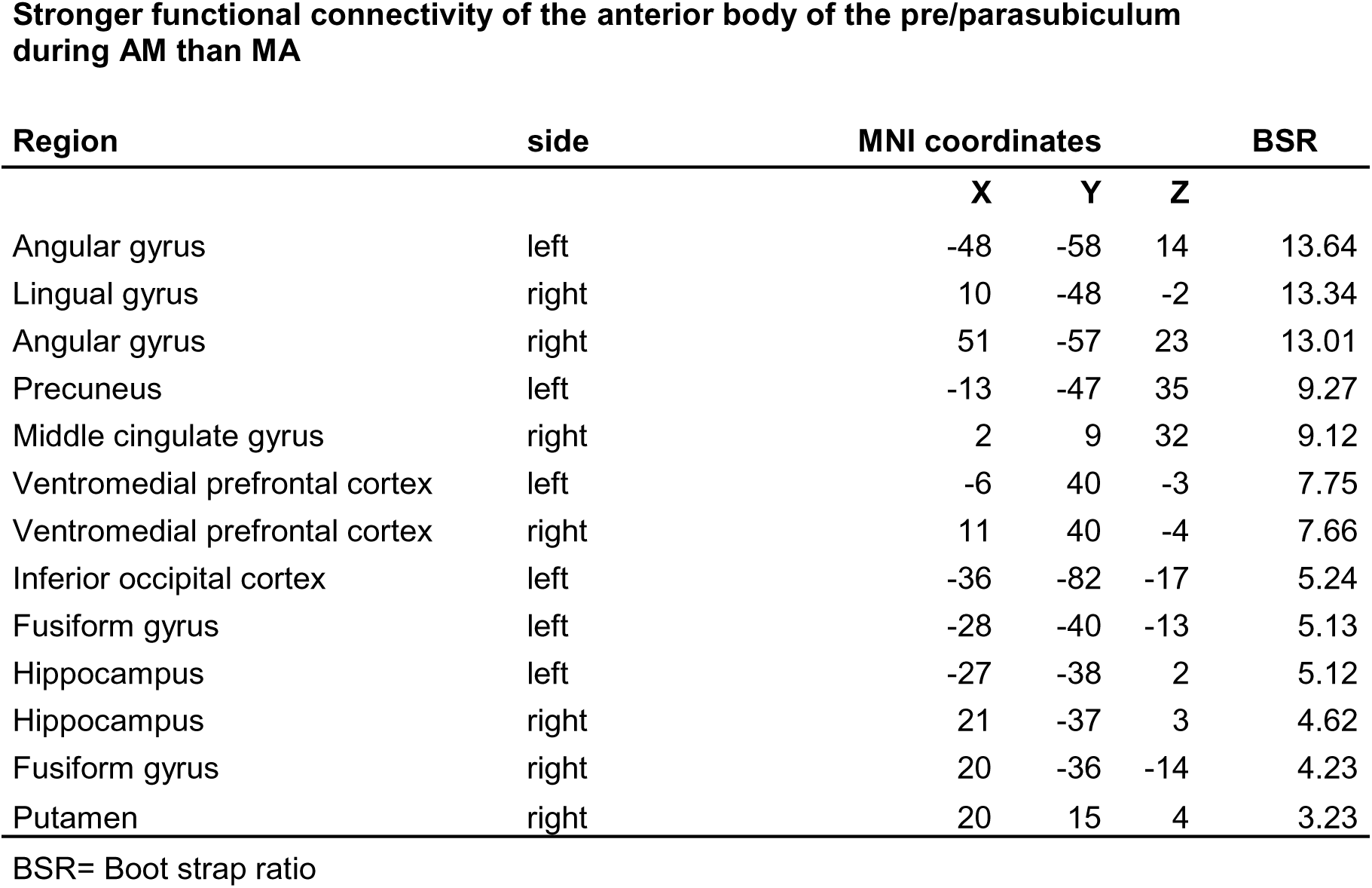
Peak coordinates of the seed PLS.

**Stronger functional connectivity of the anterior body of the pre/parasubiculum during AM than MA**
| Region | side | MNI coordinates |  |  | BSR |
| --- | --- | --- | --- | --- | --- |
|  |  | X | Y | Z |  |
| Angular gyrus | left | -48 | -58 | 14 | 13.64 |
| Lingual gyrus | right | 10 | -48 | -2 | 13.34 |
| Angular gyrus | right | 51 | -57 | 23 | 13.01 |
| Precuneus | left | -13 | -47 | 35 | 9.27 |
| Middle cingulate gyrus | right | 2 | 9 | 32 | 9.12 |
| Ventromedial prefrontal cortex | left | -6 | 40 | -3 | 7.75 |
| Ventromedial prefrontal cortex | right | 11 | 40 | -4 | 7.66 |
| Inferior occipital cortex | left | -36 | -82 | -17 | 5.24 |
| Fusiform gyrus | left | -28 | -40 | -13 | 5.13 |
| Hippocampus | left | -27 | -38 | 2 | 5.12 |
| Hippocampus | right | 21 | -37 | 3 | 4.62 |
| Fusiform gyrus | right | 20 | -36 | -14 | 4.23 |
| Putamen | right | 20 | 15 | 4 | 3.23 |

BSR= Boot strap ratio

**Table S5.** Peak coordinates of the seed PLS.

**Stronger functional connectivity of the anterior body of the pre/parasubiculum during MA than AM.**
| Region | side | MNI coordinates |  |  | BSR |
| --- | --- | --- | --- | --- | --- |
|  |  | X | Y | Z |  |
| Dorsolateral prefrontal cortex | left | -58 | 17 | 9 | -9.61 |
| Supplementary motor cortex | right | 6 | -17 | 54 | -7.04 |
| Cerebellum | right | 14 | -69 | -46 | -6.52 |
| Ventrolateral prefrontal cortex | left | -20 | 54 | -6 | -5.84 |
| Cerebellum | left | -13 | -79 | -45 | -4.63 |
| Temporal pole | left | -27 | 6 | -23 | -4.21 |

BSR= Boot strap ratio

## References

Addis DR, Wong AT, Schacter DL. Remembering the past and imagining the future: common and distinct neural substrates during event construction and elaboration. Neuropsychologia 2007; 45: 1363–1377.

Auger ML, Floresco SB. Prefrontal cortical GABA modulation of spatial reference and working memory. Int. J. Neuropsychopharmacol. 2014; 18

Baker S, Vieweg P, Gao F, Gilboa A, Wolbers T, Black SE, et al. The human dentate gyrus plays a necessary role in discriminating new memories. Curr. Biol. 2016; 26: 2629–2634.

Barry DN, Clark IA, Maguire EA. The relationship between hippocampal subfield volumes and autobiographical memory persistence. Hippocampus 2021; 31: 362–374.

Bartsch T, Döhring J, Rohr A, Jansen O, Deuschl G. CA1 neurons in the human hippocampus are critical for autobiographical memory, mental time travel, and autonoetic consciousness. Proceedings of the National Academy of Sciences 2011;108(42):17562–7.

Berron D, Schütze H, Maass A, Cardenas-Blanco A, Kuijf HJ, Kumaran D, et al. Strong evidence for pattern separation in human dentate gyrus. J. Neurosci. 2016; 36: 7569–7579.

Boccara CN, Sargolini F, Thoresen VH, Solstad T, Witter MP, Moser EI, et al. Grid cells in pre- and parasubiculum. Nat. Neurosci. 2010; 13: 987–994.

Bonnici HM, Chadwick MJ, Kumaran D, Hassabis D, Weiskopf N, Maguire EA. Multi-voxel pattern analysis in human hippocampal subfields. Front. Hum. Neurosci. 2012; 6: 290.

Bonnici HM, Chadwick MJ, Lutti A, Hassabis D, Weiskopf N, Maguire EA. Detecting representations of recent and remote autobiographical memories in vmPFC and hippocampus. J. Neurosci. 2012; 32: 16982–16991.

Bonnici HM, Chadwick MJ, Maguire EA. Representations of recent and remote autobiographical memories in hippocampal subfields. Hippocampus 2013;23(10):849–54.

Brenner D, Stirnberg R, Pracht ED, Stöcker T. ESMRMB 2013, 30th Annual Scientific Meeting, Toulouse, France, 3-5 October: Abstracts, Friday. Magn Reson Mater Phy 2013; 26: 151–301.

Breuer FA, Blaimer M, Mueller MF, Seiberlich N, Heidemann RM, Griswold MA, et al. Controlled aliasing in volumetric parallel imaging (2D CAIPIRINHA). Magn. Reson. Med. 2006; 55: 549–556.

Chadwick MJ, Bonnici HM, Maguire EA. CA3 size predicts the precision of memory recall. Proc. Natl. Acad. Sci. USA 2014; 111: 10720–10725.

Clark IA, Maguire EA. Do questionnaires reflect their purported cognitive functions? Cognition 2020; 195: 104114.

Conway MA, Pleydell-Pearce CW. The construction of autobiographical memories in the self-memory system. Psychol. Rev. 2000; 107: 261–288.

Conway MA. Episodic memories. Neuropsychologia 2009; 47: 2305–2313.

Coupé P, Manjón JV, Gedamu E, Arnold D, Robles M, Collins DL. Robust Rician noise estimation for MR images. Med Image Anal 2010; 14: 483–493.

Dalton MA, D’Souza A, Lv J, Calamante F. New insights into anatomical connectivity along the anterior–posterior axis of the human hippocampus using in vivo quantitative fibre tracking. Elife. 2022; 11; e76143.

Dalton MA, Maguire EA. The pre/parasubiculum: a hippocampal hub for scene-based cognition? Curr. Opin. Behav. Sci. 2017; 17: 34–40.

Dalton MA, McCormick C, De Luca F, Clark IA, Maguire EA. Functional connectivity along the anterior-posterior axis of hippocampal subfields in the ageing human brain. Hippocampus 2019; 29: 1049–1062.

Dalton MA, McCormick C, Maguire EA. Differences in functional connectivity along the anterior-posterior axis of human hippocampal subfields. Neuroimage 2019; 192: 38–51.

Dalton MA, Zeidman P, Barry DN, Williams E, Maguire EA. Segmenting subregions of the human hippocampus on structural magnetic resonance image scans: An illustrated tutorial. Brain Neurosci. Adv. 2017; 1: 2398212817701448.

Dalton MA, Zeidman P, McCormick C, Maguire EA. Differentiable Processing of Objects, Associations, and Scenes within the Hippocampus. J. Neurosci. 2018; 38: 8146–8159.

Dawes AJ, Keogh R, Andrillon T, Pearson J. A cognitive profile of multi-sensory imagery, memory and dreaming in aphantasia. Sci. Rep. 2020; 10: 10022.

Dice LR. Measures of the Amount of Ecologic Association Between Species. Ecology 1945; 26: 297.

Ding S-L, Van Hoesen GW. Organization and Detailed Parcellation of Human Hippocampal Head and Body Regions Based on a Combined Analysis of Cyto- and Chemoarchitecture. J. Comp. Neurol. 2015; 523: 2233–2253.

Ding S-L. Comparative anatomy of the prosubiculum, subiculum, presubiculum, postsubiculum, and parasubiculum in human, monkey, and rodent. J. Comp. Neurol. 2013; 521: 4145–4162.

Epstein RA, Parker WE, Feiler AM. Where am I now? Distinct roles for parahippocampal and retrosplenial cortices in place recognition. J. Neurosci. 2007; 27: 6141–6149.

Ezama L, Hernández-Cabrera JA, Seoane S, Pereda E, Janssen N. Functional connectivity of the hippocampus and its subfields in resting-state networks. Eur. J. Neurosci. 2021; 53: 3378–3393.

Finn ES, Huber L, Jangraw DC, Molfese PJ, Bandettini PA. Layer-dependent activity in human prefrontal cortex during working memory. Nat. Neurosci. 2019; 22: 1687–1695.

Grande X, Berron D, Horner AJ, Bisby JA, Düzel E, Burgess N. Holistic recollection via pattern completion involves hippocampal subfield CA3. J. Neurosci. 2019; 39: 8100–8111.

Grande X, Sauvage MM, Becke A, Düzel E, Berron D. Transversal functional connectivity and scene-specific processing in the human entorhinal-hippocampal circuitry. Elife. 2022; 11; e76479.

Greenberg DL, Knowlton BJ. The role of visual imagery in autobiographical memory. Mem. Cognit. 2014; 42: 922–934.

Hennig J, Scheffler K. Hyperechoes. Magn. Reson. Med. 2001; 46: 6–12.

Hodgetts CJ, Voets NL, Thomas AG, Clare S, Lawrence AD, Graham KS. Ultra-High-Field fMRI Reveals a Role for the Subiculum in Scene Perceptual Discrimination. J. Neurosci. 2017; 37: 3150–3159.

Kravitz DJ, Saleem KS, Baker CI, Mishkin M. A new neural framework for visuospatial processing. Nat. Rev. Neurosci. 2011; 12: 217–230.

Krishnan A, Williams LJ, McIntosh AR, Abdi H. Partial Least Squares (PLS) methods for neuroimaging: a tutorial and review. Neuroimage 2011; 56: 455–475.

Lever C, Burton S, Jeewajee A, O’Keefe J, Burgess N. Boundary vector cells in the subiculum of the hippocampal formation. J. Neurosci. 2009; 29: 9771–9777.

Maass A, Schütze H, Speck O, Yonelinas A, Tempelmann C, Heinze H-J, et al. Laminar activity in the hippocampus and entorhinal cortex related to novelty and episodic encoding. Nat. Commun. 2014; 5: 5547.

Maguire EA, Mullally SL. The hippocampus: a manifesto for change. J. Exp. Psychol. Gen. 2013; 142: 1180–1189.

McCormick C, Barry DN, Jafarian A, Barnes GR, Maguire EA. vmPFC Drives Hippocampal Processing during Autobiographical Memory Recall Regardless of Remoteness. Cereb. Cortex 2020; 30: 5972–5987.

McCormick C, Ciaramelli E, De Luca F, Maguire EA. Comparing and contrasting the cognitive effects of hippocampal and ventromedial prefrontal cortex damage: A review of human lesion studies. Neuroscience 2018; 374: 295–318.

McCormick C, Dalton MA, Zeidman P, Maguire EA. Characterising the hippocampal response to perception, construction and complexity. Cortex 2021; 137: 1–17.

McCormick C, Rosenthal CR, Miller TD, Maguire EA. Hippocampal damage increases deontological responses during moral decision making. J. Neurosci. 2016; 36: 12157–12167.

McCormick C, Rosenthal CR, Miller TD, Maguire EA. Deciding what is possible and impossible following hippocampal damage in humans. Hippocampus 2017; 27: 303–314.

McCormick C, Rosenthal CR, Miller TD, Maguire EA. Mind-Wandering in People with Hippocampal Damage. J. Neurosci. 2018; 38: 2745–2754.

McCormick C, St-Laurent M, Ty A, Valiante TA, McAndrews MP. Functional and effective hippocampal-neocortical connectivity during construction and elaboration of autobiographical memory retrieval. Cereb. Cortex 2015; 25: 1297–1305.

McIntosh AR, Lobaugh NJ. Partial least squares analysis of neuroimaging data: applications and advances. Neuroimage 2004; 23 Suppl 1: S250–63.

Miller TD, Chong TT-J, Aimola Davies AM, Johnson MR, Irani SR, Husain M, et al. Human hippocampal CA3 damage disrupts both recent and remote episodic memories. Elife 2020; 9

Milton F, Fulford J, Dance C, Gaddum J, Heuerman-Williamson B, Jones K, et al. Behavioral and Neural Signatures of Visual Imagery Vividness Extremes: Aphantasia versus Hyperphantasia. Cereb. Cortex Commun. 2021; 2: tgab035.

Monzel M, Vetterlein A, Reuter M. Memory deficits in aphantasics are not restricted to autobiographical memory - Perspectives from the Dual Coding Approach. J. Neuropsychol. 2021

Moscovitch M, Rosenbaum RS, Gilboa A, Addis DR, Westmacott R, Grady C, et al. Functional neuroanatomy of remote episodic, semantic and spatial memory: a unified account based on multiple trace theory. J. Anat. 2005; 207: 35–66.

Newman EL, Hasselmo ME. CA3 sees the big picture while dentate gyrus splits hairs. Neuron 2014; 81: 226–228.

Norris DG, Polimeni JR. Laminar (f)MRI: A short history and future prospects. Neuroimage 2019; 197: 643–649.

Palombo DJ, Amaral RSC, Olsen RK, Müller DJ, Todd RM, Anderson AK, et al. KIBRA polymorphism is associated with individual differences in hippocampal subregions: evidence from anatomical segmentation using high-resolution MRI. J. Neurosci. 2013; 33: 13088–13093.

Palombo DJ, Bacopulos A, Amaral RSC, Olsen RK, Todd RM, Anderson AK, et al. Episodic autobiographical memory is associated with variation in the size of hippocampal subregions. Hippocampus 2018; 28: 69–75.

Palombo DJ, Sheldon S, Levine B. Individual differences in autobiographical memory. Trends Cogn. Sci. (Regul. Ed.) 2018; 22: 583–597.

Pearson J. The human imagination: the cognitive neuroscience of visual mental imagery. Nat. Rev. Neurosci. 2019; 20: 624–634.

Poppenk J, Evensmoen HR, Moscovitch M, Nadel L. Long-axis specialization of the human hippocampus. Trends in cognitive sciences 2013;17(5):230–40.

Risius U-M, Staniloiu A, Piefke M, Maderwald S, Schulte FP, Brand M, et al. Retrieval, monitoring, and control processes: a 7 tesla FMRI approach to memory accuracy. Front. Behav. Neurosci. 2013; 7: 24.

Robertson RG, Rolls ET, Georges-François P, Panzeri S. Head direction cells in the primate pre-subiculum. Hippocampus 1999; 9: 206–219.

Robin J, Buchsbaum BR, Moscovitch M. The primacy of spatial context in the neural representation of events. J. Neurosci. 2018; 38: 2755–2765.

Rosenbaum RS, Moscovitch M, Foster JK, Schnyer DM, Gao F, Kovacevic N, et al. Patterns of autobiographical memory loss in medial-temporal lobe amnesic patients. J. Cogn. Neurosci. 2008; 20: 1490–1506.

Scoville WB, Milner B. Loss of recent memory after bilateral hippocampal lesions. J. Neurol. Neurosurg. Psychiatry 1957; 20: 11–21.

Sekeres MJ, Winocur G, Moscovitch M. The hippocampus and related neocortical structures in memory transformation. Neurosci. Lett. 2018; 680: 39–53.

Shah P, Bassett DS, Wisse LEM, Detre JA, Stein JM, Yushkevich PA, et al. Mapping the structural and functional network architecture of the medial temporal lobe using 7T MRI. Hum. Brain Mapp. 2018; 39: 851–865.

Shine JP, Valdés-Herrera JP, Tempelmann C, Wolbers T. Evidence for allocentric boundary and goal direction information in the human entorhinal cortex and subiculum. Nat. Commun. 2019; 10: 4004.

Stirnberg R, Brenner D, Stöcker T, Shah NJ. Rapid fat suppression for three-dimensional echo planar imaging with minimized specific absorption rate. Magn. Reson. Med. 2016; 76: 1517– 1523.

Stirnberg R, Huijbers W, Brenner D, Poser BA, Breteler M, Stöcker T. Rapid whole-brain resting-state fMRI at 3 T: Efficiency-optimized three-dimensional EPI versus repetition time-matched simultaneous-multi-slice EPI. Neuroimage 2017; 163: 81–92.

Stirnberg R, Stöcker T. Segmented K-space blipped-controlled aliasing in parallel imaging for high spatiotemporal resolution EPI. Magn. Reson. Med. 2021; 85: 1540–1551.

Strange BA, Witter MP, Lein ES, Moser EI. Functional organization of the hippocampal longitudinal axis. Nature Reviews Neuroscience. 2014;15(10):655–69.

Svoboda E, McKinnon MC, Levine B. The functional neuroanatomy of autobiographical memory: A meta-analysis. Neuropsychologia 2006; 44: 2189–2208.

van der Kouwe AJW, Benner T, Salat DH, Fischl B. Brain morphometry with multiecho MPRAGE. Neuroimage 2008; 40: 559–569.

van Dijk MT, Fenton AA. On how the dentate gyrus contributes to memory discrimination. Neuron 2018; 98: 832–845.e5.

Watkins NW. (A)phantasia and severely deficient autobiographical memory: Scientific and personal perspectives. Cortex 2018; 105: 41–52.

Willems T, Henke K. Imaging human engrams using 7 Tesla magnetic resonance imaging. Hippocampus 2021; 31: 1257–1270.

Yoo, PE, John, SE, Farquharson, S, Cleary, JO, Wong, YT, Ng, A, Mulcahy, CB, et al. 7T-fMRI: Faster temporal resolution yields optimal BOLD sensitivity for functional network imaging specifically at high spatial resolution. Neuroimage 2018; 164; 214–229.

Yushkevich PA, Piven J, Hazlett HC, Smith RG, Ho S, Gee JC, et al. User-guided 3D active contour segmentation of anatomical structures: significantly improved efficiency and reliability. Neuroimage 2006; 31: 1116–1128.

Zeidman P, Lutti A, Maguire EA. Investigating the functions of subregions within anterior hippocampus. Cortex 2015; 73: 240–256.

Zeidman P, Maguire EA. Anterior hippocampus: the anatomy of perception, imagination and episodic memory. Nat. Rev. Neurosci. 2016; 17: 173–182.

Zeman A, Dewar M, Della Sala S. Lives without imagery - Congenital aphantasia. Cortex 2015; 73: 378–380.

Zeman A, Milton F, Della Sala S, Dewar M, Frayling T, Gaddum J, et al. Phantasia-The psychological significance of lifelong visual imagery vividness extremes. Cortex 2020; 130: 426– 440.

